# Mitochondrial substitution rates estimation for molecular clock analyses in modern birds based on full mitochondrial genomes

**DOI:** 10.1101/855833

**Authors:** Angel Arcones, Raquel Ponti, David R. Vieites

## Abstract

Abstractti

Mitochondrial DNA (mtDNA) is a very popular resource in the study of evolutionary processes in birds, and especially to infer divergence times between lineages. These inferences rely on rates of substitution in the mtDNA genes that, ideally, are specific for the studied taxa. But as such values are often unavailable many studies fixed rate values generalised from other studies, such as the popular “standard molecular clock”. However the validity of these universal rates across all bird lineages and for the different mtDNA has been severely questioned. Thus, we here performed the most comprehensive calibration of the mtDNA molecular clock in birds, with the inclusion of complete mitochondrial genomes for 622 bird species and 25 reliable fossil calibrations. The results show variation in the rates between lineages and especially between markers, contradicting the universality of the standard clock. Moreover, we provide especific rates for every mtDNA marker (except D-loop) in each of the sampled avian orders, which should help improve future estimations of divergence times between bird species or populations.

## Introduction

Mitochondrial DNA (mtDNA) is a useful and accessible source of genetic data to explore the recent evolutionary histories of species. Its extended use has been the cornerstone of many phylogeographic studies on bird species (Avise & Walker, 1998; Zink & Barrowclough, 2008), although it has also raised concerns around the reliability of specific methodologies (Ballard & Whitlock, 2004; Ho, 2007; Edwards & Bensch, 2009).

One of the most popular uses of mtDNA in ornithology is the estimation of divergence times. Given the poor fossil record for many avian families and orders (Mayr, 2009), mtDNA mutation rates have been primarily used to estimatedivergence times between lineages via molecular clock analyses. However, a large proportion of the studies based their estimations on the use of a “universal” standard molecular clock. This popular rate proposes a fixed value of 2% divergence per million years between any two lineages, which translates into approximately 0.01 substitutions per site per lineage per million years (s/s/l/My). This rate was initially estimated based on data from humans and chimpanzees (Brown *et al*., 1979) and was supported by similar rate estimations for other vertebrates (Wilson *et al*., 1985) and a couple genera of geese (Shields & Wilson, 1987). More recently, a reanalysis extended the applicability of this general rate to all birds (Weir & Schluter, 2008). This standard rate has served as a foundation of many studies involving dating divergence events on birds, such as the study of the intra and interspecific diversification during the Pleistocene (Klicka & Zink, 1997; Avise & Walker 1998; Johnson & Cicero, 2004, Weir & Schluter, 2004).

Despite of its widespread use, the validity of this standard molecular clock for any mtDNA loci in any bird lineage has been heavily criticized (Garcia-Moreno, 2004; Lovette, 2004; Pereira & Baker, 2006; Ho, 2007). Previous analyses of the mtDNA of birds highlight a lack of “standard” rates, finding variation between the different mtDNA genes as well as across different avian lineages (Pereira & Baker, 2006; Pacheco *et al*., 2011; Lavinia *et al*., 2016; Nabholz *et al*., 2016; Nguyen & Ho, 2016). Some of these studies also attempted to calibrate the molecular rates of the different protein-coding mitochondrial genes across the avian phylogeny using complete mitochondrial genomes (Pereira & Baker, 2006; Pacheco *et al*., 2011; Nabholz *et al*., 2016). However, these studies show high variation between their results, and more importantly, they often fail to provide rates that are specific for each mtDNA marker in each lineage of the tree. This is of particular importance as proper divergence time estimations rely on using substitution rates as specific as possible in each study case (Garcia-Moreno, 2004; Ho *et al*., 2007, 2015).

To overcome this problem, we here performed a comprehensive mitogenomic calibration of the avian molecular clock to obtain substitution rates for each mtDNA gene across the phylogenetic tree of avian species. We hope that these new rates will help future evolutionary and phylogeographic studies on birds to improve their molecular clock dating analyses and better understand the evolutionary processes within the different taxa of interest.

## Materials and methods

### Complete mitochondrial genomes

We assembled a comprehensive dataset of complete bird mitochondrial genomes available from Genbank by November 2016. To retrieve the genomes, we modified the Mitobank script (Abascal *et al*., 2007) in BioPerl, providing complete or nearly complete bird genomes for 622 species representing 33 modern out of 38 bird orders, including some extinct species. Of these 622 species sampled 17 belonged to palaeognaths (Ostrich, tinamous, and allies), 94 to the Galloanserae clade (chickens, quails, geese, fowls and allies) and 511 to the Neoaves (all other avian groups), of which 259 were Passeriformes (Accession Numbers: see Supplementary material). Only five orders were left unrepresented due to the lack of species’ mt genome available data: Mesitornithiformes (Mesites), Pterocliformes (Sandgrouses and allies), Opisthocomiformes (Hoatzin), Leptosomiformes (Cuckoo rollers) and Cariamiformes (Seriemas and allies).

In two cases where the original genomes lack one locus, those were completed with sequence data from other individuals of the same species: *Psittacus erithacus* for ND6 (accession number KM611474) and *Accipiter gularis* for ND1 (EU583261).

After excluding the D-loop and the tRNAs, the result was a combined dataset of 12950 base pairs. Alignments were done gene by gene as global alignments are not possible because of gene rearrangements and complexity of the genomes (Mueller & Boore, 2005). The alignment was performed using the software Genious v.9 (https://www.geneious.com). Due to the high variability of some regions of the mitochondrial DNA, and especially in 3rd codon positions, we applied a translation alignment. This implies translating the sequences into aminoacids for the alignment and then returning to the original nucleotide sequence. The genes were then concatenated into a single sequence per species.

We computed a phylogenetic tree using RaxML (Stamatakis, 2014) as a test to detect troublesome sequences that could be introducing errors in the process. This resulted in the exclusion of *Larus vegae* (GenBank accession number: NC_029383) from further analyses, as the sequence does not belong to a gull.

### Fossil time constraints

In order to accurately determine divergence times and evolutionary rates, analyses must include many independent fossil constraints to time-calibrate the phylogenetic tree (Near & Sanderson, 2004; Benton & Donoghue, 2007; Magallón *et al*., 2013). In our case, these calibrations are based on the fossil record. Following recent recommendations (Ho & Duchêne, 2014; Zheng & Wiens, 2015), we aimed to include the higher number of calibrations as possible across the tree to maximize the accuracy of our results. We included 25 calibration constraints based on their informative value and compatibility with our dataset. Following the recommendations from Parham *et al*. (2012) and Fourment & Holmes (2014), we used a conservative minimum age based on the chronostratigraphic evidence of each fossil, and we did not impose any hard-maximum ages. The calibrated nodes with their corresponding minimum ages are: (1) split *Corvus* – *Urocisa*, 13.6 Mya (Scofield *et al*., 2017); (2) split Psittaciformes – Passeriformes, 53.5 Mya (Ksepka & Clarke, 2015); (3) crown Melegridinae + Tetraoninae, 16 Mya ptarmigan (Stein *et al*., 2015); (4) crown Galliformes, 51.6 Mya(Mayr & Weidig, 2004); (5) Crown Cracide + Numididae + Phasianidae, 33.9 Mya (van Tuinen & Dyke, 2004); (6) crown *Gallus* + *Coturnix*, 23.03 Mya (van Tuinen & Dyke, 2004); (7) Crown Anatidae, 66.5 Mya (Ksepka & Clarke, 2015); (8) split Apodidae – Trochilidae, 53 Mya (Harrison, 1984); (9) crown Trochilidae, 30 Mya (Mayr, 2004); (10) split Charadriiformes – sister taxa, 46.5 Mya (Mayr, 2000); (11) crown Laromorphae, 23.6 Mya (Smith, 2015); (12) split Jacanidae – other Scolopacids, 30 Mya (Smith, 2015); (13) split Pan-Alcidae – Stercorariidae, 34.2 (Smith, 2015); (14) crown Calidrinae, 11.62 (Smith, 2015); (15) split Phoenicopteriformes – Podicipediformes, 27.8 Mya (Mayr & Smith, 2002); (16) split Fregatidae – Suloidea, 51.8 Mya (Smith & Ksepka, 2015); (17) split Phalacrocoracidae – Anhingidae, 24.5 Mya (Smith & Ksepka, 2015); (18) crown Balaenicipitidae, 30 Mya (Smith, 2013); (19) split Sphenisciformes – Procellariiformes, 60.5 Mya (Ksepka & Clarke, 2015); (20) split *Spheniscus* – *Eudyptula*, 11 Mya (Göhlich, 2007); (21) crown Uppupidae + Phoeniculidae, 46.6 Mya (Mayr, 2006; Ksepka & Clarke, 2015); (22) split Trogoniformes – sister taxa, 49 Mya (Mayr, 2005); (23) split Falconidae – Polynorinae, 15.9 Mya (Becker, 1987); (24) crown Columbiformes, 20.3 Mya (Gradstein & Ogg, 2004); (25) split *Casuarius* – *Dromaius*, 23 Mya (Boles, 1992).

### Rate estimates

We performed Bayesian analyses as implemented in BEAST2 v.2.4.7 (Bouckaert *et al*., 2014) to estimate the substitution rate per locus. The choice of this software was based on its capability to accommodate different parameters into the analysis (such as substitution models for each codon position and multiple options of relaxed molecular clock), as well as the capability of recovering a rate for each gene in all the nodes of the tree. We set independent partitions for every gene to estimate the molecular clock rate per locus, and for every codon position to fit the nucleotide substitution models selected. We used PartitionFinder (Lanfear *et al*., 2012) to estimate the best substitution model for each codon position in each gene.

We selected the lognormal relaxed clock implemented in BEAST2. In order to avoid misleading results due to clock selection, we also ran an analysis under a random local clock model to compare between results. Due to the fast rate of change in mtDNA, saturation becomes an issue affecting the topology at deeper nodes. To minimize this problem, we constrained the tree topology by grouping species’ sequences into their corresponding orders, as well as the phylogenetic relationships between orders based on the results from Prum *et al*. (2015). Relationships between Galliformes families were also constrained to fit that topology. The phylogenetic tree was computed using all genes combined, following a birth-death Yule model.

Given the nature of the dataset, with an extensive taxon sampling and basepairs, the MCMC chain needed a large number of generations in order to reach stability. We ran the chain for 1 billion (1×10^9^) generations during several months, discarding the first 500 million as burnin and sampling every 5000 states. To confirm that the previous results were not stuck on a local maximum, we ran a second 6×10^8^ independent chain to compare with. The random local clock analysis also ran for 5×10^8^ generations. Additionally, we ran an analysis using only the information from the priors, to determine whether the posterior probability of the results is driven solely by the priors, or if the data are also introducing information into the final results.

We also ran an analysis based on aminoacids and using the same specifications as in the nucleotide analyses, with substitution modelMTREV for each of the partitioned markers as implemented in BEAST2. Aminoacid sequences are more conserved than nucleotide sequences, allowing an assessment of the potential effect of mtDNA saturation of nucleotides in the topology and divergence time estimates.

## Results

### Molecular rate estimates

BEAST analyses provided a fully resolved tree topology with most nodes recovered as highly supported (>98% of nodes above 0.9 posterior probability). The Maximum Clade Credibility method implemented in TreeAnnotator forcefully prevents polytomies, but no signs of conflict were found in the resolution of any of the nodes of the tree. Results from the sample from prior differed from those from the main analyses, indicating that the recovered posterior probabilities are not just a product of the priors. The random clock analysis recovered a phylogeny where most of the nodes of the tree were younger than 1 Mya. Thus, we considered this method unreliable and discarded it as an alternative. The topology of the amino acid tree (not shown) provided similar results to the obtained from nucleotides, except minor differences at the position of particular tips, affecting less than 5% of the species.

The resulting phylogenetic tree had overall high support, with over 98% of the nodes showing a posterior probability above 0.9. We recovered a root age (Palaeognatha / Neognatha split) of 88.3Mya (95% CI = 90.9 – 85.4 Mya). The split between Neoaves and Galloanserae was recovered at 87.9 Mya (90.7 – 85.3 Mya). The divergence between all extant avian orders occurred before or around the transition between the Cretaceous and the Paleogene, 66Mya, and most of their diversification took place during the Paleogene and Neogene.

In general, the fossil-calibrated nodes returned ages several million years older than the prior minimum age of the fossils. The only two exceptions were the calibrations for the Sphenisciformes / Procellariiformes split (minimum age = 60.5Mya) and the *Dromaius* / *Casuarius* split (min. age = 23.03Mya). In both cases, the posterior distribution was constrained by the hard minimum bound of the prior, but still displayed a normal distribution and the mean ages were slightly older than the minimum constraint (not shown).

Our approach allowed us to obtain substitution rates for each gene across each node of the tree. We found variation in the rates between genes, and to a lesser extent, between lineages in the tree in each gene. The highly-conserved ribosomal genes 12S and 16S showed the lowest mean substitution rate over the whole tree (0.00107 and 0.00043 s/s/l/My, respectively), while ND2, ATP6, and ATP8 returned the highest overall rate values (0.00227, 0.00215 and 0.00260 s/s/l/My, respectively) (Fig. 1).

**Figure 1:**
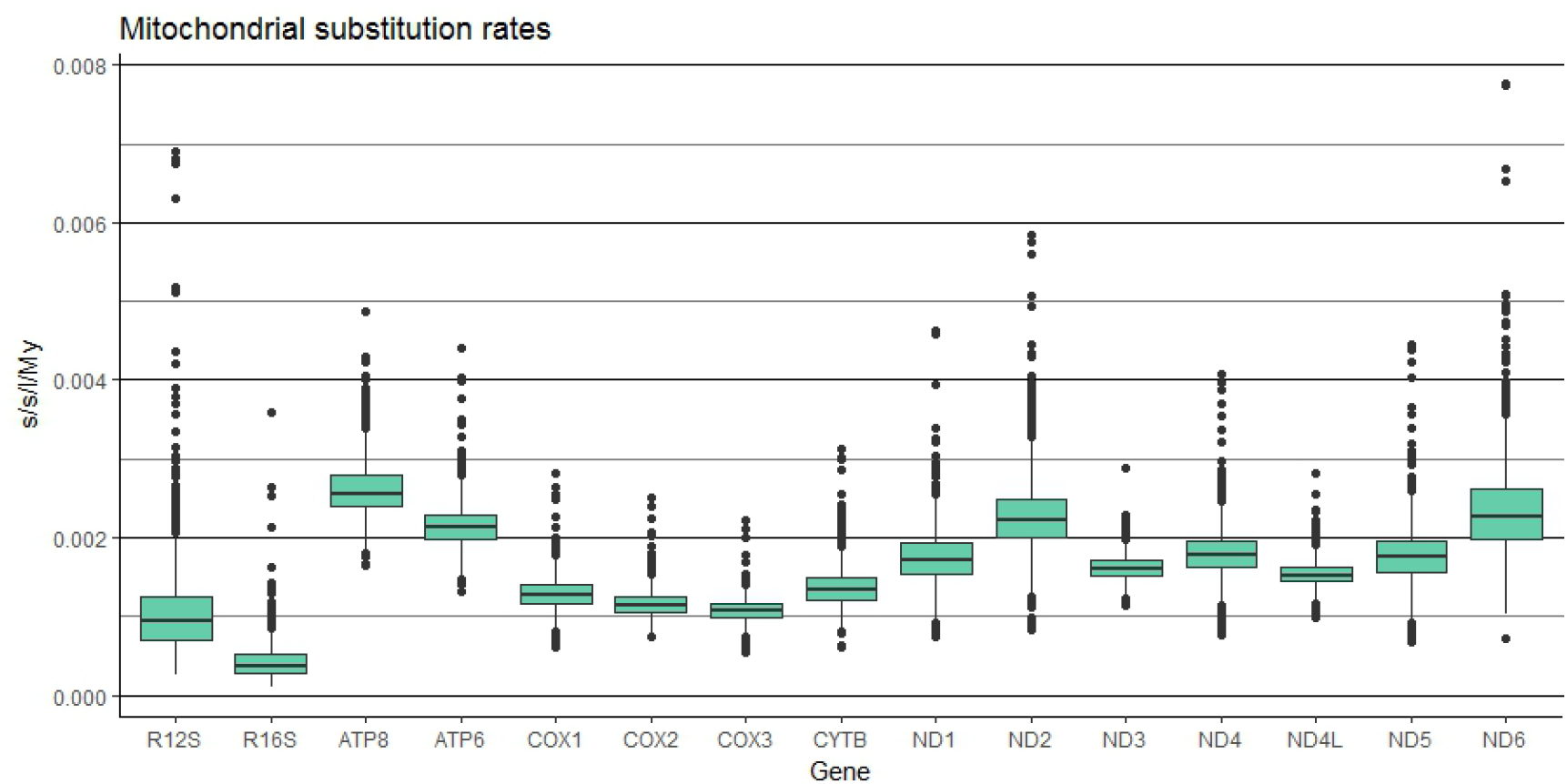
boxplot of the substitution rate values (in substitutions / site / lineage / million years) from every node of the mitogenomic tree, for each of the 15 analyzed mitochondrial genes.

Overall, the rate values in each gene showed minimal variation across the branches of the tree, and there were no significant differences in the overall mitochondrial rates between orders (Fig. 2). Only the order Otidiformes showed values distributed mostly below the overall average, although this order is represented by just a single species in the sampling (*Otis tarda*), and could be affected by a punctual underestimation of the rates due to insufficient taxon coverage. The values of the rates per gene for the avian orders sampled can be found in the Supplementary material.

**Figure 2:**
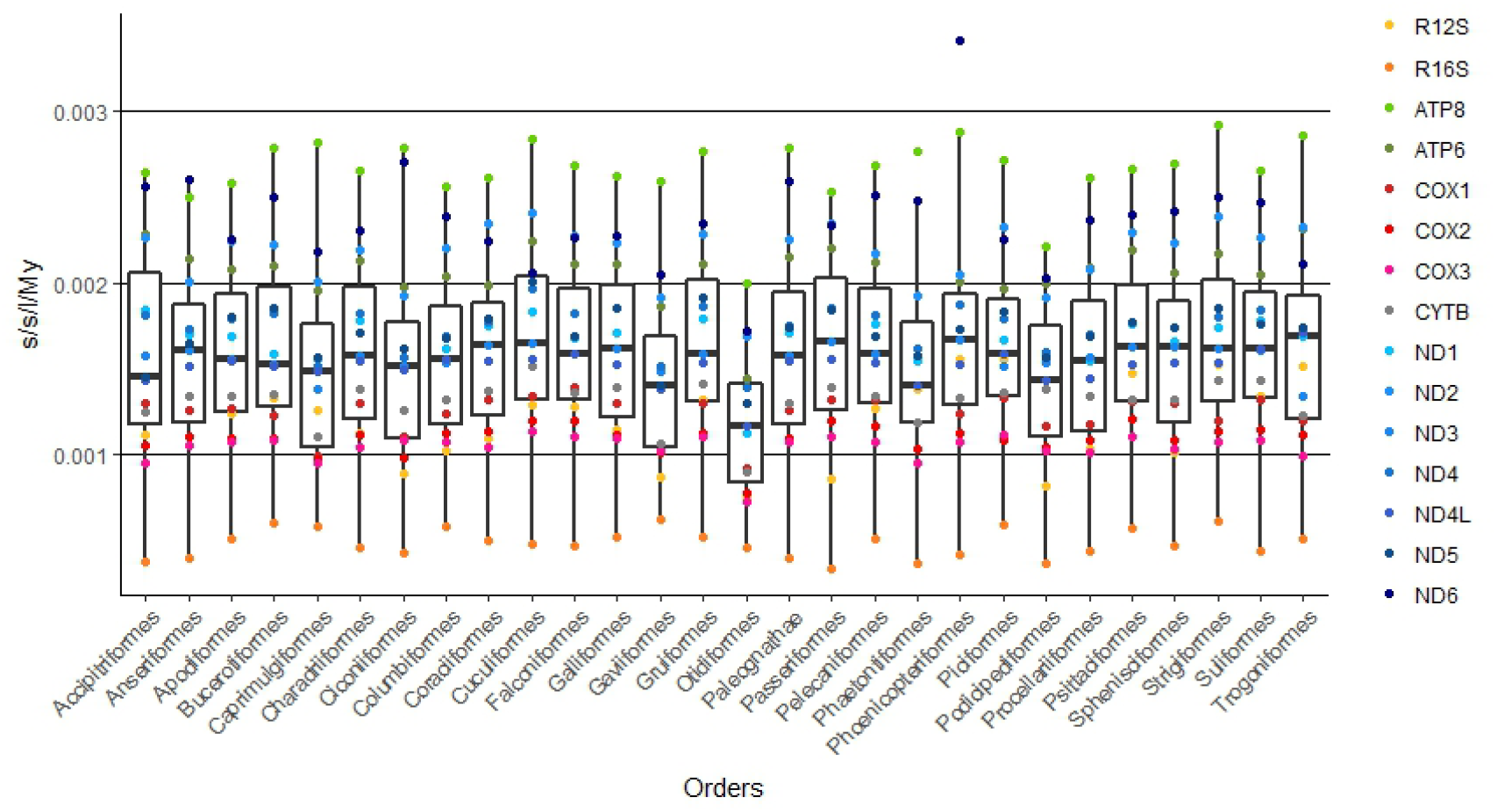
boxplot of the average substitution rate values (in substitutions / site / lineage / million years) of all 15 mitochondrial genes for each of the orders of birds.

## Discussion

Our analyses represent the most comprehensive calibration of the mitochondrial molecular clock performed to date in birds, both in terms of species coverage and fossil calibrations used. We provide the first lineage-specific rates for each of the mtDNA genes for almost all of the extant bird orders (Fig. 2, Suplementary material).

The results from previous mitogenomic studies (Pereira & Baker, 2006; Pacheco *et al*., 2011; Nabholz *et al*., 2016) showed significant variability in the recovered evolutionary rates. In all cases they found rate values lower than the so-called standard molecular clock of 0.01 s/s/l/My (Fig 3). The substitution rate values obtained by us fall within the range of the variation observed between previous works (Fig. 3). Moreover, the differences in mean rate values for the mitochondrial genes in our study follow a similar pattern, although with lower value than the rates obtained by Nabholz *et al*. (2016) (Fig. 3, blue line), which until now was the most taxonomic-rich and phylogenetically updated analysis.

**Figure 3:**
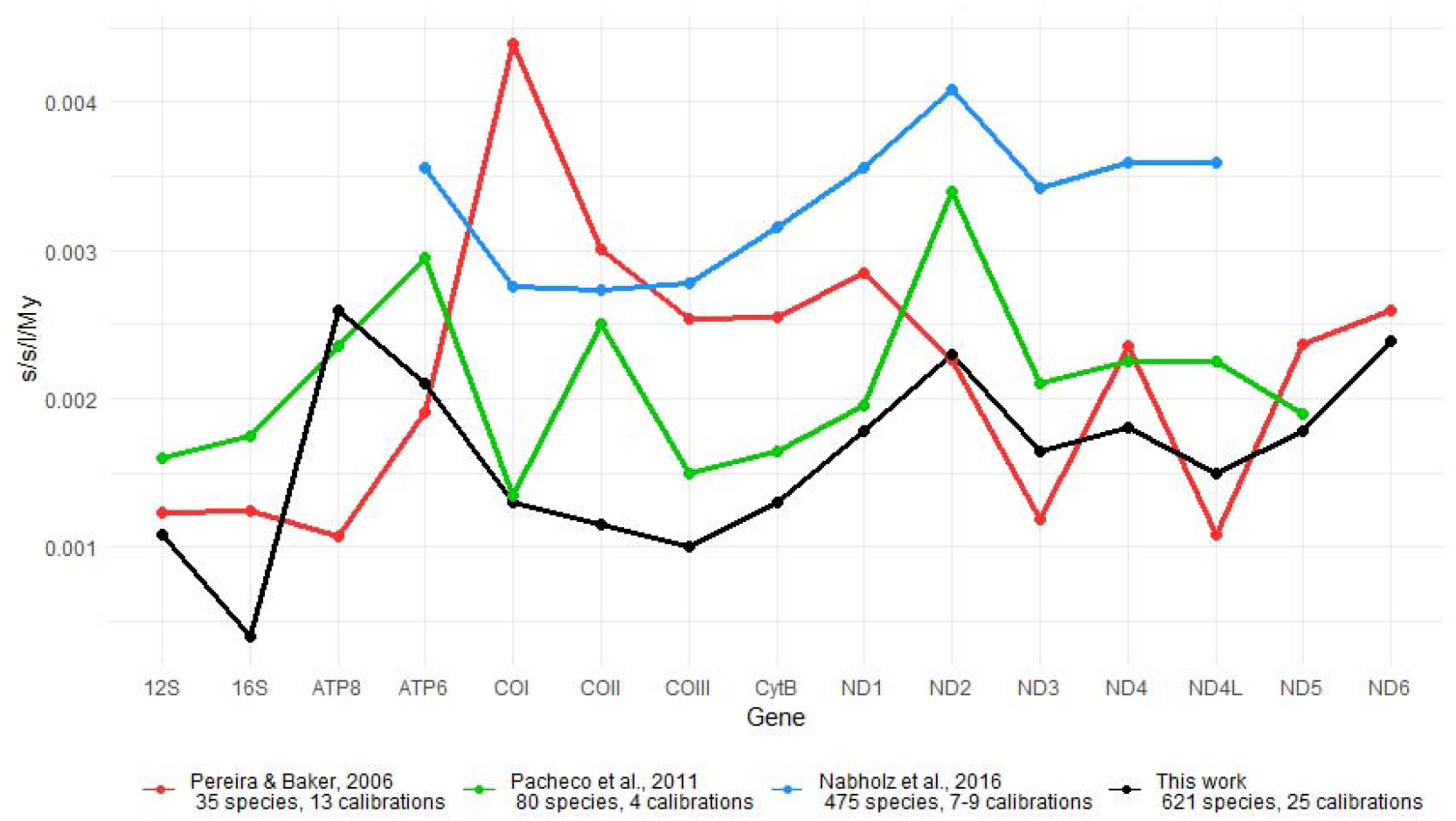
Comparison of the mean substitution rate estimates for each mitochondrial gene, between our results (black line) and previous mitogenomic calibrations of the molecular clock, with the number of species and fossil time constraints included in each analysis.

The differences in mutation rates between studies can be the consequence of several causes, from the inclusion or elimination of third codon positions that are generally saturated at deeper phylogenetic levels, to the impact of hard maximum bounds in fossil time constraints that force younger ages in the phylogeny (Ho *et al*., 2005, 2007, 2008), and even the choice of software or other specific parameters could be introducing variation (Drummond & Rambaut, 2015). Given the variety of methodologies that have been used in previous mitogenomic rate estimations for birds (Pereira & Baker, 2006; Pacheco *et al*., 2011; Nabholz *et al*., 2016), we are unable to assess whether the differences between our results and those from previous studies are caused by one or several of the mentioned causes, or by any other cause. In any case, we are confident that given the more extensive taxonomic coverage, the more and reliable fossil calibrations included, and the methodological approach (e.g. no maximum-time constraints, very long Bayesian MCMC runs until reaching true stabilization), our results provide a reliable estimation of the molecular clock substitution rates in birds.

Our results, along with those from previous analyses, contradict the validity of the standard mitochondrial molecular clock (Pereira & Baker, 2006; Pacheco *et al*., 2011; Nabholz *et al*., 2016) and support the criticism against its generalized use (Garcia-Moreno, 2004; Lovette, 2004; Ho, 2007, Lavinia *et al*., 2016). These new substitution rates per locus provided in this study should improve the accuracy and reliability of future molecular clock analyses, making available rates for more clades that can be used instead of standard rates. We expect that the implementation of reliable well-calibrated rates will significantly benefit future research on the recent evolutionary history of bird groups without fossils to time-calibrate their genealogy orphylogenetic relationships.

## Supporting information

Supplementary 1: Genbank Accesion Numbers

Supplementary 2: Substitution rates for each order

