## Supplementary 1: Genbank Accesion Numbers for "Mitochondrial substitution rates estimation for molecular clock analyses in modern birds based on full mitochondrial genomes"

**Supplementary material 1: Genbank accession numbers for the species' mitochondrial genomes.**

| Species | Accesion number |
|---|---|
| *Abrornis inornata(Phylloscopus)* | NC_024726 |
| *Acanthis flammea* | NC_027285 |
| *Acanthisitta chloris* | AY325307 |
| *Accipiter gentilis* | NC_011818 |
| *Accipiter gularis* | KX585864 / EU583261 |
| *Accipiter nisus* | NC_025580 |
| *Accipiter soloensis* | KJ680303 |
| *Accipiter virgatus* | NC_026082 |
| *Aceros corrugatus* | HM755883 |
| *Aceros waldeni* | NC_015085 |
| *Acridotheres cristatellus* | NC_015613 |
| *Acridotheres tristis (Sturnus)* | NC_015195 |
| *Acrocephalus scirpaceus* | NC_010227 |
| *Acryllium vulturinum* | NC_014180 |
| *Aegithalos bonvaloti* | NC_024267 |
| *Aegithalos caudatus* | KF951088 |
| *Aegithalos concinnus* | KF951092 |
| *Aegithalos fuliginosus* | NC_024266 |
| *Aegithalos glaucogularis* | NC_024268 |
| *Aegotheles cristatus* | NC_011718 |
| *Aegypius monachus* | KF682364 |
| *Aepyornis hildebrandti* | KJ749824 |
| *Aethopyga gouldiae* | NC_027241 |
| *Agapornis roseicollis* | NC_011708 |
| *Agelaius phoeniceus* | NC_018801 |
| *Aix galericulata* | NC_023969 |
| *Akialoa obscura* | NC_031349 |
| *Alauda arvensis* | NC_020425 |
| *Alectoris chukar* | NC_020585 |
| *Alectura lathami* | NC_007227 |
| *Amaurornis akool* | NC_023982 |
| *Amaurornis phoenicurus* | NC_024593 |
| *Amazilia versicolor* | NC_024156 |
| *Amazona barbadensis* | JX524615 |
| *Amazona ochrocephala* | NC_027840 |
| *Amblyramphus holosericeus* | NC_018802 |
| *Anas acuta* | NC_024631 |
| *Anas chathamica* | KF562761 |
| *Anas clypeata* | NC_028346 |
| *Anas crecca* | NC_022452 |
| *Anas formosa* | NC_015482 |
| *Anas platyrhynchos* | NC_009684 |
| *Anas poecilorhyncha* | NC_022418 |
| *Anhinga rufa* | GU071055 |
| *Anomalopteryx didiformis* | NC_002779 |
| *Anser albifrons* | NC_004539 |
| *Anser anser* | NC_011196 |
| *Anser cygnoides* | NC_023832 |
| *Anser fabalis* | NC_016922 |
| *Anser indicus* | NC_025654 |
| *Anseranas semipalmata* | NC_005933 |
| *Anthornis melanura* | KC545408 |
| *Anthropoides paradiseus* | NC_020572 |
| *Anthropoides virgo* | NC_020573 |
| *Anthus hodgsoni* | KX189345 |
| *Anthus novaeseelandiae* | NC_029137 |
| *Aphrodroma brevirostris* | NC_007174 |
| *Aptenodytes forsteri* | NC_027938 |
| *Apteryx australis* | KU695537 |
| *Apteryx haastii* | NC_002782 |
| *Apteryx owenii* | NC_013806 |
| *Apus apus* | NC_008540 |
| *Aquila chrysaetos* | NC_024087 |
| *Ara ararauna* | NC_029319 |
| *Ara glaucogularis* | NC_026029 |
| *Ara militaris* | NC_027839 |
| *Aratinga (Psittaccara) mitrata* | JX215256 |
| *Arborophila ardens* | NC_022683 |
| *Arborophila brunneopectus* | NC_022684 |
| *Arborophila gingica* | FJ752425 |
| *Arborophila rufipectus* | NC_012453 |
| *Arborophila rufogularis* | NC_020584 |
| *Archilochus colubris* | NC_010094 |
| *Ardea cinerea* | NC_025900 |
| *Ardea intermedia* | NC_025918 |
| *Ardea modesta* | NC_025916 |
| *Ardea novaehollandiae* | NC_008551 |
| *Ardea purpurea* | NC_025919 |
| *Ardeola bacchus* | NC_025921 |
| *Arenaria interpres* | NC_003712 |
| *Argusianus argus* | JQ713768 |
| *Arremon aurantiirostris* | NC_027731 |

| | |
|---|---|
| *Asio flammeus* | NC_027606 |
| *Athene brama* | KF961185 |
| *Aythya americana* | NC_000877 |
| *Aythya ferina* | NC_024602 |
| *Aythya fuligula* | NC_024595 |
| *Babax lanceolatus* | KR818090 |
| *Balaeniceps rex* | GU071053 |
| *Balearica pavonina* | NC_020570 |
| *Balearica regulorum* | NC_020569 |
| *Bambusicola fytchii* | NC_020583 |
| *Bambusicola thoracica* | NC_011816 |
| *Bombycilla cedrorum* | KJ909187 |
| *Botaurus stellaris* | NC_025923 |
| *Branta bernicla* | KJ680301 |
| *Branta canadensis* | NC_007011 |
| *Brotogeris cyanoptera* | NC_015530 |
| *Bubo blakistoni* | LC099103 |
| *Bubo bubo* | AB918148 |
| *Bubo flavipes* | LC099100 |
| *Bubulcus ibis* | NC_025917 |
| *Bucorvus leadbeateri* | NC_015199 |
| *Buteo buteo* | NC_003128 |
| *Buteo hemilasius* | NC_029377 |
| *Buteo lagopus* | NC_029189 |
| *Butorides striata* | NC_025922 |
| *Bycanistes brevis* | NC_015201 |
| *Cacatua moluccensis* | NC_020592 |
| *Cacatua pastinator* | JF414240 |
| *Cairina moschata* | NC_010965 |
| *Callaeas cinereus* | NC_031350 |
| *Callipepla squamata* | NC_029340 |
| *Calliphlox amethystina* | NC_030286 |
| *Caloperdix oculeus* | NC_024619 |
| *Calyptorhynchus baudinii* | NC_020594 |
| *Calyptorhynchus lathami* | NC_020593 |
| *Calyptorhynchus latirostris* | NC_020595 |
| *Campephilus guatemalensis* | NC_028020 |
| *Campephilus imperialis* | KU158198 |
| *Campylorhynchus brunneicapillus* | NC_029482 |
| *Campylorhynchus zonatus* | NC_022840 |
| *Caprimulgus indicus* | NC_025773 |
| *Cardinalis cardinalis* | NC_025618 |
| *Carduelis (Spinus) psaltria* | NC_025627 |
| *Carduelis (Spinus) spinus* | NC_015198 |
| *Carduelis pinus* | NC_025619 |
| *Carduelis sinica* | NC_015196 |
| *Carpodacus erythrinus* | NC_025597 |
| *Carpodacus roseus* | NC_025607 |
| *Casuarius casuarius* | NC_002778 |
| *Cathartes aura* | NC_007628 |
| *Cecropis daurica* | NC_024107 |
| *Centropus sinensis* | KT947122 |
| *Ceryle rudis* | NC_024280 |
| *Chaetura pelagica* | NC_028545 |
| *Chalcophaps indica* | HM746789 |
| *Chlorophanes spiza* | NC_025606 |
| *Chroicocephalus ridibundus* | NC_025649 |
| *Chrysolampis mosquitus* | NC_025786 |
| *Chrysolophus amherstiae* | NC_020590 |
| *Chrysolophus pictus* | NC_014576 |
| *Chrysomus cyanopus* | NC_018813 |
| *Chrysomus icterocephalus* | NC_018799 |
| *Chrysomus ruficapillus* | NC_018796 |
| *Chrysomus thilius* | NC_018807 |
| *Chrysomus xanthophthalmus* | NC_018798 |
| *Ciconia boyciana* | NC_002196 |
| *Ciconia ciconia* | NC_002197 |
| *Ciconia nigra* | NC_023946 |
| *Cnemotriccus fuscatus* | NC_007975 |
| *Coccothraustes coccothraustes* | NC_025614 |
| *Colinus virginianus* | NC_024620 |
| *Columba janthina* | KM926619 |
| *Columba livia* | NC_013978 |
| *Copsychus saularis* | NC_030603 |
| *Coracopsis vasa* | NC_027841 |
| *Corvus brachyrhynchos* | NC_026461 |
| *Corvus cornix* | NC_024698 |
| *Corvus frugilegus* | NC_002069 |
| *Corvus hawaiiensis* | NC_026783 |
| *Corvus macrorhynchos* | NC_027173 |
| *Corvus moriorum* | NC_031518 |
| *Corvus splendens* | NC_024607 |
| *Coturnicops exquisitus* | NC_012143 |
| *Coturnix chinensis* | NC_004575 |
| *Coturnix japonica* | NC_003408 |
| *Crax daubentoni* | NC_024617 |
| *Crax rubra* | NC_024618 |
| *Crossoptilon auritum* | NC_015897 |
| *Crossoptilon crossoptilon* | NC_016679 |
| *Crossoptilon harmani* | NC_026547 |
| *Crossoptilon mantchuricum* | NC_026548 |
| *Crotophaga ani* | HM746794 |

| | |
|---|---|
| *Cuculus poliocephalus* | NC_028414 |
| *Curaeus curaeus* | NC_018808 |
| *Cyanistes cyanus* | KX388472 |
| *Cyanopica cyanus* | NC_015824 |
| *Cyanoptila cyanomelana* | NC_015232 |
| *Cygnus atratus* | NC_012843 |
| *Cygnus columbianus* | NC_017604 |
| *Cygnus cygnus* | NC_027095 |
| *Cygnus olor* | NC_027096 |
| *Dendrocopos leucotos* | NC_029862 |
| *Dendrocopos major* | NC_028174 |
| *Dendrocygna javanica* | NC_012844 |
| *Deroptyus accipitrinus* | KM611476 |
| *Dinornis giganteus* | NC_002672 |
| *Diomedea chrysostoma* | AP009193 |
| *Dives dives* | NC_018800 |
| *Dromaius novaehollandiae* | NC_002784 |
| *Dryocopus pileatus* | NC_008546 |
| *Dupetor flavicollis* | NC_024575 |
| *Eclectus roratus* | NC_027842 |
| *Ectopistes migratorius* | KC489473 |
| *Egretta eulophotes* | NC_009736 |
| *Egretta garzetta* | NC_023981 |
| *Egretta sacra* | NC_025920 |
| *Emberiza (Schoeniclus) elegans* | NC_030368 |
| *Emberiza (Schoeniclus) rustica* | NC_024924 |
| *Emberiza (Schoeniclus) spodocephala* | NC_021445 |
| *Emberiza aureola (Schoeniclus aureolus)* | NC_022150 |
| *Emberiza chrysophrys* | NC_015233 |
| *Emberiza cioides* | NC_024524 |
| *Emberiza jankowskii* | NC_027251 |
| *Emberiza pusilla* | NC_021408 |
| *Emberiza rutila* | NC_024925 |
| *Emberiza tristrami* | NC_015234 |
| *Emeus crassus* | NC_002673 |
| *Eophona migratoria* | NC_031374 |
| *Eopsaltria australis* | NC_019665 |
| *Eopsaltria georgiana* | NC_027230 |
| *Eopsaltria griseogularis* | NC_027229 |
| *Epthianura albifrons* | NC_019664 |
| *Eremopsaltria mongolica* | NC_025616 |
| *Eudocimus ruber* | NC_027504 |
| *Eudromia elegans* | NC_002772 |
| *Eudynamys taitensis* | NC_011709 |
| *Eudyptes chrysocome* | NC_008138 |

| | |
|---|---|
| *Eudyptula minor* | NC_004538 |
| *Eulabeornis castaneoventris* | NC_025501 |
| *Euphagus cyanocephalus* | NC_018827 |
| *Eupsittula pertinax* | NC_015197 |
| *Eurynorhynchus pygmeus* | NC_027496 |
| *Eurystomus orientalis* | NC_011716 |
| *Falco cherrug* | NC_026715 |
| *Falco columbarius* | NC_025579 |
| *Falco naumanni* | NC_029846 |
| *Falco peregrinus* | NC_000878 |
| *Falco rusticolus* | NC_029359 |
| *Falco sparverius* | NC_008547 |
| *Falco tinnunculus* | NC_011307 |
| *Ficedula zanthopygia* | NC_015802 |
| *Fidecula albicollis* | NC_021621 |
| *Florisuga fusca* | NC_030287 |
| *Florisuga mellivora* | NC_027455 |
| *Forpus modestus* | HM755882 |
| *Forpus passerinus* | NC_027843 |
| *Francolinus pintadeanus* | NC_011817 |
| *Fregata sp.* | AP009192 |
| *Fringilla coelebs* | NC_025599 |
| *Fringilla montifringilla* | NC_024048 |
| *Fringilla polatzeki* | NC_031157 |
| *Fringilla teydea* | KU705760 |
| *Fulica atra* | NC_025500 |
| *Gallicolumba luzonica* | HM746790 |
| *Gallicrex cinerea* | NC_028408 |
| *Gallinula chloropus* | NC_015236 |
| *Gallirallus australis* | KF425525 |
| *Gallirallus okinawae* | NC_012140 |
| *Gallirallus philippensis* | NC_025507 |
| *Gallus gallus* | NC_001323 |
| *Gallus lafayetii* | NC_007239 |
| *Gallus sonneratii* | NC_007240 |
| *Gallus varius* | NC_007238 |
| *Garrulax affinis* | NC_029402 |
| *Garrulax canorus* | NC_020429 |
| *Garrulax cineraceus* | NC_024553 |
| *Garrulax ocellatus* | NC_027657 |
| *Garrulax perspicillatus* | NC_026068 |
| *Garrulax poecilorhynchus* | NC_028082 |
| *Garrulax sannio* | NC_028186 |
| *Garrulus glandarius* | NC_015810 |
| *Gavia pacifica* | NC_008139 |
| *Gavia stellata* | NC_007007 |

| | |
|---|---|
| *Geococcyx californianus* | NC_011711 |
| *Geopelia striata* | HM746791 |
| *Geospiza fortis* | KM891730 |
| *Geotrygon violacea* | NC_015207 |
| *Gerygone igata* | NC_029139 |
| *Glaucidium brodiei* | KP684122 |
| *Gnorimopsar chopi* | NC_018795 |
| *Gorsachius goisagi* | NC_028194 |
| *Gorsachius magnificus* | NC_028193 |
| *Gorsachius melanolophus* | NC_028195 |
| *Goura cristata* | LN589994 |
| *Goura scheepmakeri* | NC_027947 |
| *Goura victoria* | LN589993 |
| *Gracula religiosa* | NC_015898 |
| *Grus americana* | NC_020576 |
| *Grus antigone* | NC_020581 |
| *Grus canadensis* | NC_020582 |
| *Grus carunculatus* | NC_020571 |
| *Grus grus* | NC_020577 |
| *Grus japonensis* | NC_020575 |
| *Grus leucogeranus* | NC_020574 |
| *Grus monacha* | NC_020578 |
| *Grus nigricollis* | NC_020579 |
| *Grus rubicunda* | NC_020580 |
| *Grus vipio* | NC_021368 |
| *Gymnomystax mexicanus* | NC_018812 |
| *Haematopus ater* | NC_003713 |
| *Haemorhous cassinii* | NC_025613 |
| *Haemorhous mexicanus* | NC_025610 |
| *Halcyon coromanda* | NC_028177 |
| *Halcyon pileata* | NC_024198 |
| *Halcyon smyrnensis* | KT965614 |
| *Heliodoxa aurescens* | NC_030285 |
| *Heliornis fulica* | NC_025499 |
| *Hemignathus flavus* | NC_025608 |
| *Hemignathus parvus* | NC_025622 |
| *Hemignathus stejnegeri* | NC_025624 |
| *Hemignathus virens* | KM078788 |
| *Hemiphaga novaeseelandiae* | NC_013244 |
| *Henicorhina leucosticta* | NC_024673 |
| *Hesperiphona vespertina* | NC_025600 |
| *Heteralocha acutirostris* | NC_031351 |
| *Hieraaetus fasciatus* | NC_029188 |
| *Himatione sanguinea* | NC_025602 |
| *Hirundo rustica* | KX398931 |
| *Hyliota flavigaster* | NC_024868 |

| | |
|---|---|
| *Hylocharis cyanus* | NC_027453 |
| *Ichthyaetus relictus* | NC_023777 |
| *Icterus mesomelas* | JX516068 |
| *Ithaginis cruentus* | NC_018033 |
| *Ixobrychus cinnamomeus* | NC_015077 |
| *Ixobrychus eurhythmus* | NC_025924 |
| *Ixobrychus sinensis* | NC_025925 |
| *Jacana jacana* | NC_024069 |
| *Jacana spinosa* | NC_024068 |
| *Lampropsar tanagrinus* | JX516057 |
| *Lanius cristatus* | NC_028333 |
| *Lanius isabellinus* | NC_027655 |
| *Lanius schach* | NC_030604 |
| *Lanius sphenocercus* | KU884610 |
| *Lanius tephronotus* | NC_021105 |
| *Larus brunnicephalus* | NC_018548 |
| *Larus crassirostris* | NC_025556 |
| *Larus dominicanus* | NC_007006 |
| *Larus vegae* | NC_029383 |
| *Leiothrix argentauris* | NC_015114 |
| *Leiothrix lutea* | NC_020427 |
| *Lepidothrix coronata* | KJ909196 |
| *Leptotila verreauxi* | NC_015190 |
| *Leucocarbo (Phalacrocorax) chalconotus* | GU071054 |
| *Leucosticte arctoa* | NC_025615 |
| *Leucosticte brandti* | NC_025604 |
| *Lewinia muelleri* | NC_025502 |
| *Lonchura punctulata* | NC_028036 |
| *Lonchura striata* | NC_029475 |
| *Lophophanes dichrous* | KX388477 |
| *Lophophorus lhuysii* | NC_013979 |
| *Lophophorus sclateri* | NC_020589 |
| *Lophura ignita* | NC_010781 |
| *Lophura nycthemera* | NC_012895 |
| *Lophura swinhoii* | NC_023779 |
| *Loxia curvirostra* | NC_025623 |
| *Loxops caeruleirostris* | NC_025605 |
| *Loxops coccineus* | NC_025612 |
| *Loxops mana* | NC_025598 |
| *Luscinia calliope* | NC_015074 |
| *Luscinia cyanura* | NC_026067 |
| *Lyrurus tetrix* | NC_024554 |
| *Machlolophus spilonotus* | KX388476 |
| *Macroagelaius imthurni* | NC_018810 |
| *Malurus melanocephalus* | NC_024873 |
| *Mareca (Anas) falcata* | NC_023352 |

| | |
|---|---|
| *Megalurus pryeri* | NC_029151 |
| *Megalurus punctatus* | NC_029138 |
| *Melamprosops phaeosoma* | NC_025617 |
| *Meleagris gallopavo* | NC_010195 |
| *Meleagris ocellata* | KU094576 |
| *Melophus lathami* | KX702277 |
| *Melopsittacus undulatus* | NC_009134 |
| *Menura novaehollandiae* | NC_007883 |
| *Mergus squamatus* | NC_016723 |
| *Micrastur gilvicollis* | NC_008548 |
| *Minla ignotincta* | NC_030588 |
| *Mionectes oleagineus* | NC_024682 |
| *Moho braccatus* | NC_031348 |
| *Mohoua novaeseelandiae* | KC545409 |
| *Molothrus aeneus* | NC_018806 |
| *Molothrus badius* | NC_018811 |
| *Montifringilla adamsi* | NC_025913 |
| *Montifringilla nivalis* | NC_025911 |
| *Montifringilla ruficollis* | NC_022815 |
| *Montifringilla taczanowskii* | NC_025914 |
| *Morus serrator* | GU071056 |
| *Motacilla alba* | NC_029229 |
| *Motacilla cinerea* | NC_027933 |
| *Motacilla lugens* | NC_029703 |
| *Mullerornis agilis* | KJ749825 |
| *Myadestes myadestinus* | NC_031352 |
| *Myiopsitta monachus* | NC_027844 |
| *Nannopterum brasilianus* | NC_029758 |
| *Neophema chrysogaster* | NC_019804 |
| *Nesopsar nigerrimus* | NC_018794 |
| *Nestor notabilis* | NC_027845 |
| *Netta rufina* | NC_024922 |
| *Ninox novaeseelandiae* | NC_005932 |
| *Ninox scutulata* | NC_029384 |
| *Nipponia nippon* | NC_008132 |
| *Nisaetus alboniger* | NC_007599 |
| *Nisaetus nipalensis* | NC_007598 |
| *Notiomystis cincta* | NC_029140 |
| *Nucifraga columbiana* | NC_022839 |
| *Numenius phaeopus* | NC_030507 |
| *Numida meleagris* | NC_006382 |
| *Nyctibius grandis* | EU344977 |
| *Nyctibius griseus* | HM746792 |
| *Nycticorax nycticorax* | NC_015807 |
| *Nymphicus hollandicus* | NC_015192 |
| *Oedistoma iliolophum (iliolophus)* | NC_024865 |

| | |
|---|---|
| *Oreomystis bairdi* | NC_025628 |
| *Oreopsar bolivianus* | NC_018797 |
| *Oreotrochilus melanogaster* | NC_027454 |
| *Oriolus chinensis* | NC_020424 |
| *Orthopsittaca manilata* | NC_029161 |
| *Otis tarda* | NC_014046 |
| *Otus bakkamoena* | NC_028163 |
| *Otus scops* | NC_028162 |
| *Pachyplichas yaldwyni* | KX369036 |
| *Padda oryzivora* | NC_028441 |
| *Pandion haliaetus* | NC_008550 |
| *Paradoxornis fulvifrons* | NC_028436 |
| *Paradoxornis nipalensis* | NC_028437 |
| *Paradoxornis webbianus* | NC_024539 |
| *Paroreomyza montana* | NC_025601 |
| *Parus major* | NC_026293 |
| *Parus monticolus* | NC_028187 |
| *Parus venustulus* | NC_026701 |
| *Passer ammodendri* | NC_029344 |
| *Passer domesticus* | NC_025611 |
| *Passer montanus* | NC_024821 |
| *Pavo cristatus* | NC_024533 |
| *Pavo muticus* | NC_012897 |
| *Pelagodroma marina* | KC875856 |
| *Pelecanus conspicillatus* | DQ780883 |
| *Penelopides panini* | NC_015087 |
| *Perdix dauurica* | NC_020588 |
| *Perdix hodgsoniae* | NC_023940 |
| *Pericrocotus ethologus* | NC_024257 |
| *Periparus ater* | NC_026223 |
| *Petroica australis* | NC_029141 |
| *Petroica boodang* | NC_019666 |
| *Petroica goodenovii* | NC_019667 |
| *Petroica macrocephala* | NC_029142 |
| *Petroica phoenicea* | NC_019668 |
| *Phaethon lepturus* | NC_027275 |
| *Phaethon rubricauda* | NC_007979 |
| *Phaethornis hispidus* | KP853098 |
| *Phaethornis malaris* | NC_030288 |
| *Phalacrocorax carbo* | NC_027267 |
| *Phalcoboenus australis* | KP064202 |
| *Phasianus colchicus* | NC_015526 |
| *Phasianus versicolor* | NC_010778 |
| *Philesturnus carunculatus* | NC_029143 |
| *Phodilus badius* | NC_023787 |
| *Phoebastria albatrus* | NC_026190 |

| | |
|---|---|
| *Phoebastria immutabilis* | NC_026189 |
| *Phoebastria nigripes* | NC_026188 |
| *Phoenicopterus roseus* | NC_010089 |
| *Phoenicopterus ruber* | NC_027934 |
| *Phoenicurus auroreus* | NC_026066 |
| *Pica pica* | NC_015200 |
| *Picathartes gymnocephalus* | KJ909200 |
| *Picoides pubescens* | NC_027936 |
| *Pinguinus impennis* | NC_031347 |
| *Pinicola enucleator* | NC_025609 |
| *Pipile pipile* | KU221053 |
| *Pitta nympha* | NC_027067 |
| *Platalea leucorodia* | NC_012772 |
| *Platalea minor* | NC_010962 |
| *Podiceps cristatus* | NC_008140 |
| *Podoces hendersoni* | NC_014879 |
| *Poecile atricapilla* | NC_024867 |
| *Poecile montanus* | KX388479 |
| *Poecile palustris* | NC_026911 |
| *Polyplectron bicalcaratum* | NC_012900 |
| *Polyplectron germaini* | NC_023264 |
| *Polyplectron napoleonis* | NC_024615 |
| *Pomatorhinus ruficollis* | NC_029769 |
| *Poospiza cabanisi* | NC_028038 |
| *Poospiza lateralis* | NC_028039 |
| *Poospiza thoracica* | NC_028037 |
| *Porphyrio hochstetteri* | NC_010092 |
| *Porphyrio porphyrio* | NC_025508 |
| *Primolius couloni* | NC_025742 |
| *Primolius maracana* | NC_029322 |
| *Prioniturus luconensis* | NC_027846 |
| *Procellaria cinerea* | AP009191 |
| *Progne chalybea* | NC_020605 |
| *Prosthemadera novaeseelandiae* | NC_029144 |
| *Prunella montanella* | NC_027284 |
| *Prunella strophiata* | KU975800 |
| *Psephotellus pulcherrimus* | NC_031358 |
| *Pseudoleistes guirahuro* | NC_018809 |
| *Pseudoleistes virescens* | NC_018805 |
| *Pseudonestor xanthophrys* | NC_025630 |
| *Pseudopodoces humilis* | NC_014341 |
| *Psittacara acuticaudatus* | NC_020325 |
| *Psittacara brevipes* | NC_021764 |
| *Psittacara rubritorquis* | NC_026042 |
| *Psittacus erithacus* | NC_027847 / KM611474 |

| | |
|---|---|
| *Psittirostra psittacea* | NC_031353 |
| *Psittrichas fulgidus* | NC_027848 |
| *Pterocnemia pennata* | NC_002783 |
| *Pteroglossus azara* | NC_008549 |
| *Ptilopachus petrosus* | NC_024616 |
| *Pucrasia macrolopha* | NC_020587 |
| *Pycnonotus melanicterus* | NC_024730 |
| *Pycnonotus sinensis* | NC_013838 |
| *Pycnonotus taivanus* | NC_013483 |
| *Pycnonotus xanthorrhous* | KX129905 |
| *Pygoscelis adeliae* | NC_021137 |
| *Pygoscelis antarcticus* | NC_021474 |
| *Pyrgilauda blanfordi* | NC_025912 |
| *Pyrgilauda davidiana* | NC_025915 |
| *Pyrrhocorax graculus* | NC_025927 |
| *Pyrrhocorax pyrrhocorax* | NC_025926 |
| *Pyrrhula pyrrhula* | NC_025625 |
| *Pyrrhura rupicola* | NC_028404 |
| *Quiscalus quiscula* | NC_018803 |
| *Rallina eurizonoides* | NC_012142 |
| *Recurvirostra avosetta* | NC_027420 |
| *Regulus calendula* | NC_024866 |
| *Regulus regulus* | NC_029837 |
| *Remiz consobrinus* | NC_021641 |
| *Rhea americana* | NC_000846 |
| *Rhipidura fuliginosa* | NC_029145 |
| *Rhynchopsitta terrisi* | NC_021771 |
| *Rhynchortyx cinctus* | KJ914547 |
| *Rhynochetos jubatus* | NC_010091 |
| *Sagittarius serpentarius* | NC_023788 |
| *Sasia ochracea* | NC_028019 |
| *Saundersilarus saundersi* | NC_017601 |
| *Sceloglaux albifacies* | KX098448 |
| *Scolopax rusticola* | NC_025521 |
| *Scytalopus magellanicus* | KJ909189 |
| *Serinus albogularis* | NC_025595 |
| *Serinus canaria* | NC_023375 |
| *Serinus dorsostriatus* | NC_025621 |
| *Sitta carolinensis* | NC_024870 |
| *Smithornis sharpei* | NC_000879 |
| *Spheniscus demersus* | NC_022817 |
| *Spilornis cheela* | NC_015887 |
| *Spizixos semitorques* | NC_029321 |
| *Stachyris ruficeps* | NC_030771 |
| *Stercorarius maccormicki* | NC_026125 |
| *Sternula albifrons* | NC_028176 |

| | |
|---|---|
| *Streptopelia chinensis* | NC_026459 |
| *Streptopelia decaocto* | KX372273 |
| *Streptopelia orientalis* | NC_031447 |
| *Strigops habroptilus* | NC_005931 |
| *Strix leptogrammica* | KC953095 |
| *Struthio camelus* | NC_002785 |
| *Sturnus cineraceus* | NC_015237 |
| *Sturnus nigricollis* | NC_020423 |
| *Sturnus sericeus* | NC_014455 |
| *Sturnus vulgaris* | NC_029360 |
| *Sylvia atricapilla* | NC_010228 |
| *Sylvia crassirostris* | NC_010229 |
| *Sylviparus modestus* | NC_026793 |
| *Synthliboramphus antiquus* | NC_007978 |
| *Synthliboramphus wumizusume* | NC_029328 |
| *Syrmaticus ellioti* | NC_010771 |
| *Syrmaticus humiae* | NC_010774 |
| *Syrmaticus reevesii* | NC_010770 |
| *Syrmaticus soemmerringi* | NC_010767 |
| *Tachybaptus novaehollandiae* | NC_010095 |
| *Tachybaptus ruficollis* | NC_024594 |
| *Tachycineta albilinea* | NC_020601 |
| *Tachycineta albiventer* | NC_020602 |
| *Tachycineta bicolor* | NC_020596 |
| *Tachycineta cyaneoviridis* | NC_020599 |
| *Tachycineta euchrysea* | NC_020598 |
| *Tachycineta leucorrhoa* | NC_020603 |
| *Tachycineta meyeni* | NC_020604 |
| *Tachycineta stolzmanni* | NC_020600 |
| *Tachycineta thalassina* | NC_020597 |
| *Tadorna ferruginea* | NC_024640 |
| *Tadorna tadorna* | NC_024750 |
| *Taeniopygia guttata* | NC_007897 |
| *Tanygnathus lucionensis* | KM611480 |
| *Tetraogallus himalayensis* | NC_027279 |
| *Tetraogallus tibetanus* | NC_023939 |
| *Tetraophasis obscurus* | NC_018034 |
| *Tetraophasis szechenyii* | NC_020613 |
| *Tetrastes bonasia* | NC_020591 |
| *Tetrastes sewerzowi* | NC_025318 |
| *Thalassarche melanophrys* | NC_007172 |

| | |
|---|---|
| *Thamnophilus nigrocinereus* | KJ909192 |
| *Thraupis episcopus* | NC_025596 |
| *Threskiornis aethiopicus* | NC_013146 |
| *Tinamus guttatus* | NC_027260 |
| *Tinamus major* | NC_002781 |
| *Todiramphus sanctus* | NC_011712 |
| *Tragopan caboti* | NC_013619 |
| *Tragopan temminckii* | NC_020586 |
| *Traversia lyalli* | KX369034 |
| *Tregellasia capito* | NC_027231 |
| *Tregellasia leucops* | NC_024871 |
| *Tringa erythropus* | NC_030585 |
| *Trogon viridis* | NC_011714 |
| *Turdus eunomus* | NC_028273 |
| *Turdus hortulorum* | NC_024552 |
| *Turdus merula* | NC_028188 |
| *Turdus migratorius* | NC_024872 |
| *Turdus naumanni* | KJ834096 |
| *Turdus philomelos* | NC_029147 |
| *Turdus rufiventris* | NC_028179 |
| *Turnagra capensis* | NC_028336 |
| *Turtur tympanistria* | HM746793 |
| *Tyto alba* | EU410491 |
| *Tyto longimembris* | KP893332 |
| *Upupa epops* | NC_028178 |
| *Uragus sibiricus* | NC_025594 |
| *Urocissa erythrorhyncha* | NC_020426 |
| *Vanellus cinereus* | NC_025514 |
| *Vanellus vanellus* | NC_025637 |
| *Vestiaria coccinea* | NC_025620 |
| *Vidua chalybeata* | NC_000880 |
| *Vireo olivaceus* | NC_024869 |
| *Xanthopsar flavus* | NC_018804 |
| *Xenicus gilviventris* | KX369033 |
| *Xenicus longipes* | KX369035 |
| *Yuhina diademata* | NC_029462 |
| *Zenaida auriculata* | NC_015203 |
| *Zoothera dauma* | KT340629 |
| *Zosterops erythropleurus* | NC_027942 |
| *Zosterops japonicus* | KT601061 |
| *Zosterops lateralis* | NC_029146 |
